# Constraints on Lanthanide Separation by Selective Biosorption

**DOI:** 10.1101/2023.10.18.562985

**Authors:** Carter Anderson, Sean Medin, James Adair, Bryce Demopoulos, Emeka Eneli, Chloe Kuelbs, Joseph Lee, Timothy J. Sheppard, Deniz Şinar, Zacharia Thurston, Mingyang Xu, Kang Zhang, Buz Barstow

## Abstract

Rare Earth Elements (REE) are essential ingredients of sustainable energy technologies^1-5^, but separation of lanthanides is considered one of the hardest problems in chemistry today^6^. Biosorption, where molecules adsorb to the surface of biological materials, offers a sustainable alternative to environmentally harmful solvent extractions currently used for REE separations. The REE-biosorption capabilities of some microorganisms already allow for REE separations that under specialized conditions are competitive with solvent extractions^7^. Our recent work has discovered the genetic basis for REE biosorption selectivity^8^. However, it is unclear if the small changes to selectivity produced by changes to single genetic loci could allow biosorption to leapfrog existing REE separation technologies. In this article we present three models of lanthanide separation by biosorption and desorption. The first model shows that if a biosorbing microbe behaves as it if has a single type of binding site with small preference for one lanthanide, then the small changes in selectivity produced by changes to a single genetic locus could reduce the length of a separation process by ≈ 25%. Large multi-locus gene edits could reduce a separation process length by almost 90%. On the other hand, if the microbe contains multiple sites each with a distinct preference for an individual lanthanide, then separations become challenging and larger genetic edits would be needed to enable high purity. However, even if these large genetic edits are not possible, high purity separations could still be possible by combining multiple microbes, each with small increases in selectivity, with the first designed to enrich for the target lanthanide, while subsequent microbes are designed to remove contaminants.

## Introduction

Rare Earth Elements (REE), typically referring to the lanthanides (lanthanum to lutetium) and sometimes scandium and yttrium, are essential ingredients for current sustainable energy technologies like high strength lightweight magnets used in electric vehicles and wind turbines^4,5^; high-efficiency lighting^3^; and battery anodes^2^; and in future sustainability technologies like lightweight high-strength alloys^1^; and room temperature superconductors^9^. As a result, the total demand for REE is projected to increase 7-fold by 2040^10^. However, the extraction of REE from ore and their subsequent purification requires high temperatures and harsh solvents, and leaves considerable quantities of toxic waste^11-13^. These processes give sustainable energy technologies reliant on REE a high environmental impact and large carbon footprint^14^.

The majority of lanthanide chemical separations utilize commercially available organic solvents and extractants^15^. The separation of a mixed lanthanide solution into individual element batches requires several extraction cycles performed with large quantities of organic solvents and expensive synthetic chelators^5,7,16^. All lanthanides exist as trivalent cations and the ionic radius difference between the largest rare earth, La^3+^, and the smallest rare earth, Lu^3+^, is only 0.17 Å (ref. 17). This means that separations of adjacent or near-adjacent lanthanides pose an enormous challenge for conventional chemical methods, requiring multiple enrichments by organic solvent extractions in extremely long mixer settler devices^18^. This results in the generation of large amounts of toxic waste. As a consequence, the United States has no capacity to produce purified lanthanides, and only two lanthanide purification plants exist outside of China^12,13,19^.

Biotechnologies could replace many of the existing methods for extracting and separating individual lanthanides. Biomining already supplies 5% and 15% of the world’s gold and copper respectively ^20^. We anticipate that a REE-biomining system will operate in three steps: (1) bioleaching metals from an ore or end-of-life feedstock like a magnet; (2) separating the lanthanides from all other metals present in the leachate (*e*.*g*., uranium and thorium from an ore like monazite, or iron from a magnet); and (3) separating individual lanthanides.

Significant progress has already been made in developing microorganisms for the first bioleaching step of the biomining process^21-23^, and in the second total lanthanide separation step^24-31^. For example, lanmodulin, a lanthanide-binding protein discovered in methylotrophic bacteria, is selective for lanthanides in the presence of molar amounts of competing metal cations^26,30^. Meanwhile, lanthanide binding tags (LBTs), attached either to the surface of *Caulobacter crescentus*^24^ or an engineered curli biofilm^27^ can selectively bind lanthanides in the presence of competing metals. Finally, both *Methylorubrum extorquens*^31^, and *E. coli* engineered with surface-displayed LBTs^28^ are able to preferentially accumulate heavy lanthanides from a mixed solution of lanthanides. Furthermore, Mattocks *et al*. recently demonstrated production of >98% pure solutions of neodymium and dysprosium in a single pass^32^ and Nelson *et al*. have created molecules with separation factors up to 213 for neodymium/dysprosium^33^. As a result of these breakthroughs, a lanthanide-bioseparation system might not need to separate individual lanthanides in the presence of high concentrations of competing metals.

Biosorption and desorption from the surface of a microbial cell offers an environmentally-friendly route for individual lanthanide-separation. Biosorption can provide high metal binding capacity at low cost^34,35^. The membranes of both gram-negative and gram-positive bacteria contain multiple sites^36^ that bind lanthanides including proteins^37^, lipids^38^, and polysaccharides^39^. Bonificio *et al*. demonstrated lanthanide-separation capability by biosorption and desorption under decreasing pH by microbes including *Roseobacter* sp. AzwK-3b and *Shewanella oneidensis* MR-1^7^. Furthermore, Bonificio *et al*. demonstrated that, under specialized conditions, lanthanides separations with *Roseobacter* sp. AzwK-3b^7^ are competitive with conventional solvent extraction. Given that no molecule is known to exist with a high separation factor for adjacent lanthanides, it is unrealistic to expect industrially-acceptable purity (*i*.*e*., 95 or 99%) in a single step biosorption and elution process. But, a separation process can (like current industrial methods) rely on repeated enrichment. The low cost of biomass (relative to purified protein) means that biosorption is uniquely well suited to this sort of enrichment process.

We hypothesize that it should be possible to engineer lanthanide-biosorbing organisms like *S. oneidensis* to optimize the preferential adsorption and desorption of individual lanthanides. If possible, this could enable even greater separation of individual lanthanides than that already achieved and for biosorption to leapfrog existing technologies. In a recent work, we characterized the genetics of lanthanide-biosorption by *S. oneidensis*. In total, we discovered 242 genes that control the overall level of REE-biosorption, and 10 that control REE-binding selectivity^8^. The functions of these genes range from synthesis of the lipopolysaccharide (LPS) layer on the outer membrane of *S. oneidensis* to assembly of mannose-sensitive hemagglutinin (MSHA) pilus. We hypothesize that the lanthanide-binding preference of *S. oneidensis* (or in fact any lanthanide-biosorbing microbe) can be altered by up- or down-regulation of genes involved in the synthesis of structural features on its outer surfaces.

However, the changes to lanthanide-biosorption selectivity that we observed were small (only 1 to 4%)^8^. If these mutant microbes with small increases in lanthanide selectivity, or further engineered microbes that combine mutations, were employed in a lanthanide separation system, would they meaningfully improve its performance? In this article, we present three theoretical models of lanthanide separation by repeated biosorption and elution and calculate the effect of small changes to lanthanide-biosorption selectivity on the length of the lanthanide-separation process. Furthermore, the mathematics used here can be easily applied to lanthanide-separation by immobilized lanthanide-chelating proteins and peptides.

## Theory

We envision a process where a mixed solution of lanthanides is repeatedly biosorbed and eluted from immobilized biosorption microbe (such as *S. oneidensis*) (**Figure 1**). The separation system consists of one or more chromatographic columns containing immobilized biomass (*e*.*g*., microbes on a filter^7^; on a biofilm on a solid support; or encapsulated in gel beads^29^). Bonificio and Clark demonstrated that two cycles of a process like this could enrich for heavy lanthanides^7^, and was competitive with conventional solvent extraction methods. A full list of all mathematical symbols used in this article is shown in **Table S1**.

**Figure 1.**
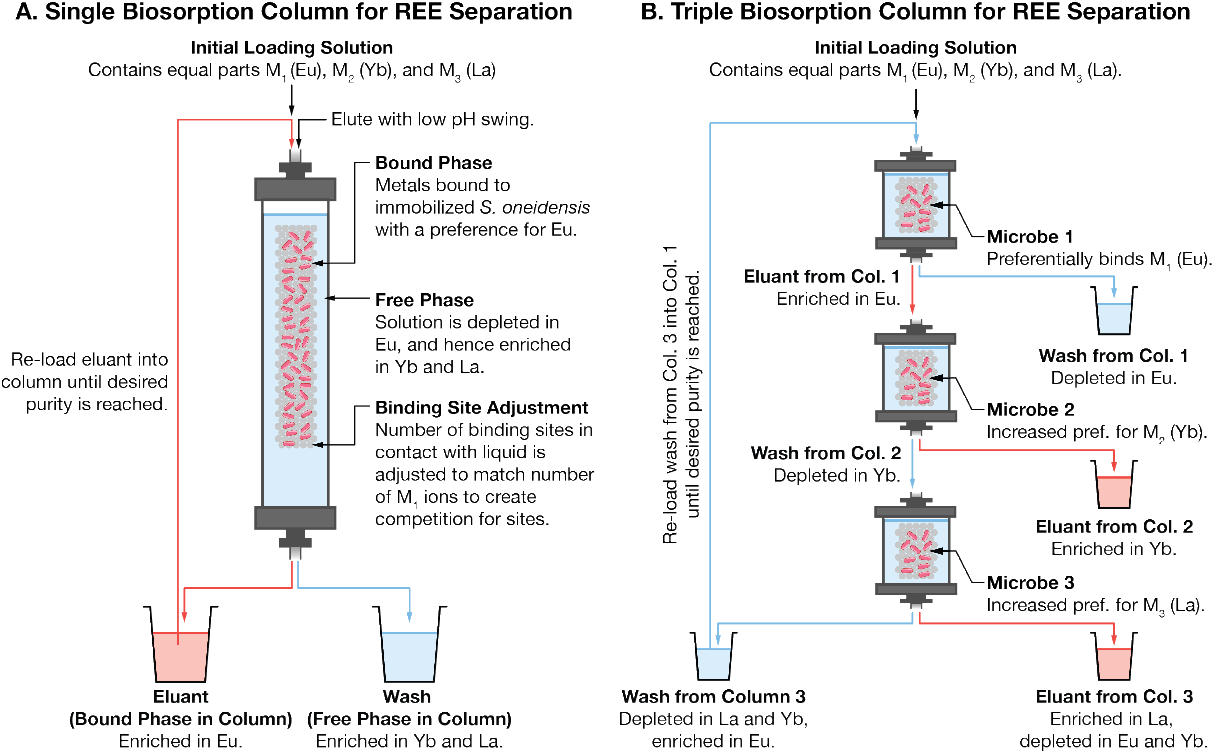
Proposed schemes for lanthanide separation by biosorption and elution. Parameters for both models are shown in **Table 1**. (**A**) Single-column scheme. The performance of this scheme for an immobilized microbe with a single type of binding site is shown in **Figure 2** and summarized in **Table 2**; and the performance with a microbe with three types of binding site is shown in **Figure 3** and summarized in **Table 3**. (**B**) Triple column scheme. We propose that this scheme should be used when the purity of the target metal (M_1_) plateaus before reaching our desired target. Performance of the three-column scheme is shown in **Figure 4**.

To sanity check the connection between experimental data and our models, we calculated the lanthanide binding per cell and per gram of dry weight of *S. oneidensis* and compared with literature values (**Note S1**, and **Table S2**). We calculate the approximate loading capacity of *S. oneidensis* is s 16.1 fg cell^-1^ / 470 fg dry weight cell^-1^ ≈ 30 mg g^-1^ of dry weight. This number compares very well to literature values of metal binding per unit mass of dry biomass assembled in **Table S2**.

**Table 1.** Model parameters used in this article.

| Parameter | Symbol | Unit | Value |
| --- | --- | --- | --- |
| **Common Parameters** |  |  |  |
| Initial number of binding sites. | $n_{B,T}$ | Mol | $3.5 \times 10^{-8}$ |
| Initial number of M<sub>1</sub> (Eu) ions | $n_{M1,T}$ | Mol | $2.4 \times 10^{-8}$ |
| Initial number of M<sub>2</sub> (Eu) ions | $n_{M2,T}$ | Mol | $2.4 \times 10^{-8}$ |
| Initial number of M<sub>3</sub> (Eu) ions | $n_{M3,T}$ | Mol | $2.4 \times 10^{-8}$ |
| Loading volume | $V_{load}$ | L | $4 \times 10^{-4}$ |
| **Single-site Model** |  |  |  |
| Dissociation constant of single site for Eu | $K_{D, Eu}$ | Mol L<sup>-1</sup> | 10<sup>-6</sup> |
| Dissociation constant of single site for Yb | $K_{D, Yb}$ | Mol L<sup>-1</sup> | $1.256 \times 10^{-6}$ |
| Dissociation constant of single site for La | $K_{D, La}$ | Mol L<sup>-1</sup> | $2.282 \times 10^{-6}$ |
| Number of binding sites | Reduces to half total number of metal ions at each step. |  |  |
| **Three-site Model** |  |  |  |
| Initial fraction of binding site 1 | $f_{B1}$ | # | 0.4618 |
| Initial fraction of binding site 2 | $f_{B2}$ | # | 0.3703 |
| Initial fraction of binding site 3 | $f_{B3}$ | # | 0.1679 |
| Dissociation constant of site 1 for Eu | $K_{D1, Eu}$ | Mol L<sup>-1</sup> | 10<sup>-6</sup> / 8 |
| Dissociation constant of site 1 for Yb | $K_{D1, Yb}$ | Mol L<sup>-1</sup> | 10<sup>-6</sup> |
| Dissociation constant of site 1 for La | $K_{D1, La}$ | Mol L<sup>-1</sup> | 10<sup>-6</sup> |
| Dissociation constant of site 2 for Eu | $K_{D2, Eu}$ | Mol L<sup>-1</sup> | 10<sup>-6</sup> |
| Dissociation constant of site 2 for Yb | $K_{D2, Yb}$ | Mol L<sup>-1</sup> | 10<sup>-6</sup> / 8 |
| Dissociation constant of site 2 for La | $K_{D2, La}$ | Mol L<sup>-1</sup> | 10<sup>-6</sup> |
| Dissociation constant of site 3 for Eu | $K_{D3, Eu}$ | Mol L<sup>-1</sup> | 10<sup>-6</sup> |
| Dissociation constant of site 3 for Yb | $K_{D3, Yb}$ | Mol L<sup>-1</sup> | 10<sup>-6</sup> |
| Dissociation constant of site 3 for La | $K_{D3, La}$ | Mol L<sup>-1</sup> | 10<sup>-6</sup> / 8 |
| Number of binding sites | Reduces to half total number of metal ions at each step. |  |  |
| **Three-site, Three-microbe Model** |  |  |  |
| Number of binding sites on microbe 1 | Reduces to total number of Eu ions at each step. |  |  |
| Number of binding sites on microbe 2 | Reduces to total number of Yb ions at each step. |  |  |
| Number of binding sites on microbe 3 | Reduces to total number of La ions at each step. |  |  |

**Table 2.** Summary of changes to REE-biosorption selectivity on REE-binding and separation with a single-site-type binding model. This table summarizes the results of **Figure 2**. (**1**) Scenario entries correspond to marked points on **Figure 2**. (**2**) Fold-decrease in the dissociation constant of the binding site for Eu. (**3**) The fraction of Eu in the metals bound to the cell surface from an equimolar solution of REE (*i*.*e*., the microbes are presented with a solution containing 33.3% Eu, 33.3% Yb, and 33.3% La, but the metals bound to the wild-type (1^st^ row) consists of 39.3% Eu. (**4**) Fractional increase of the amount of Eu in the bound phase (*f*_Eu, b_) relative to wild-type. (**5**) The separation factor between Eu and Yb. (**6** to **8**) Number of biosorption/elution cycles needed to reach 90, 95, and 99% purity of Eu. This table can be reproduced with the programs Fig-2A-B.PY, Fig-2C-D.PY, and Fig-2E.py in the ree-purification repository^41^.

| 1. Scenario | 2. Single-site $K_{D \text{ Eu}, 0} / K_{D \text{ Eu}, \text{New}}$ | 3. Initial Purity of Eu | 4. Increase in Purity of Eu in Bound Phase Over Wild-type | 5. $a_{\text{Eu}}^{\text{Yb}}$ | 6. Single-site, 90% Purity | 7. Single-site, 95% Purity | 8. Single-site, 99% Purity |
| --- | --- | --- | --- | --- | --- | --- | --- |
| 1 | 1.000 | 0.393 | 1.000 | 1.256 | 20 | 26 | 40 |
| 2 | 1.043 | 0.397 | 1.010 | 1.310 | 18 | 23 | 36 |
| 3 | 1.081 | 0.401 | 1.020 | 1.358 | 16 | 21 | 31 |
| 4 | 1.163 | 0.409 | 1.041 | 1.461 | 13 | 17 | 25 |
| 5 | 1.286 | 0.420 | 1.069 | 1.615 | 10 | 13 | 19 |
| 6 | 1.552 | 0.440 | 1.120 | 1.950 | 8 | 9 | 14 |
| 7 | 2.847 | 0.500 | 1.272 | 3.576 | 5 | 6 | 8 |
| 8 | 6.027 | 0.560 | 1.425 | 7.572 | 3 | 4 | 5 |

**Table 3.** Summary changes to REE-biosorption selectivity on REE-separation with a three-site-type binding model. This table summarizes the results of **Figure 3**. (**1**) Entries correspond to marked points on **Figure 3**. (**2**) Population-fraction of the Eu-preferring site on the surface of *S. oneidensis*. (**3**) Calculated fraction of Eu in the metals bound to the cell surface from an equimolar solution of REE (*i*.*e*., the microbes are presented with a solution containing 33.3% Eu, 33.3% Yb, and 33.3% La, but the metals bound to the wild-type (1^st^ row) consists of 39.3% Eu). (**4**) Calculated fractional increase of the amount of Eu in the bound phase. (**5**) Calculated separation factor between Eu and Yb. (**6** to **9**) Number of biosorption/elution cycles needed to reach 80, 90, 95, and 99% purity of Eu. If no number is listed then purity cannot be achieved. This table can be reproduced with the programs Fig-3A-B.PY, Fig-3C-D.PY, and Fig-3E.py in the ree-purification repository^41^.

| 1. Scenario | 2. Population-fraction of Eu-site, $f_{b, \text{Eu}}$ | 3. Initial Purity of Eu | 4. Increase in Purity of Eu in Bound Phase Over Wild-type | 5. $a_{\text{Eu}}^{\text{Yb}}$ | 6. Single-site, 80% Purity | 7. Single-site, 90% Purity | 8. Single-site, 95% Purity | 9. Single-site, 99% Purity |
| --- | --- | --- | --- | --- | --- | --- | --- | --- |
| 1 | 0.463 | 0.393 | 1.000 | 1.263 |  |  |  |  |
| 2 | 0.473 | 0.397 | 1.010 | 1.314 |  |  |  |  |
| 3 | 0.482 | 0.401 | 1.020 | 1.369 |  |  |  |  |
| 4 | 0.502 | 0.409 | 1.041 | 1.484 |  |  |  |  |
| 5 | 0.700 | 0.478 | 1.217 | 3.152 | 8 |  |  |  |
| 6 | 0.800 | 0.506 | 1.287 | 4.424 | 4 | 7 | 22 |  |
| 7 | 0.850 | 0.517 | 1.317 | 5.183 | 3 | 5 | 8 | 28 |
| 8 | 0.900 | 0.528 | 1.344 | 6.030 | 3 | 4 | 6 | 11 |
| 9 | 0.950 | 0.538 | 1.369 | 6.968 | 3 | 4 | 4 | 7 |

In our model, the system is loaded with a solution initially containing equimolar quantities of three metals (M_1_, M_2_ and M_3_; *e*.*g*., Eu, Yb and La) in a volume *V*_load_. The column contains *n*_B,T_ binding sites. After equilibration the free fraction (the liquid) is removed from the column and moved to a wash collection container. Next, the bound fraction is eluted (for example by a pH swing^7,40^). The eluant is then pH adjusted (to compensate for the pH swing), and its volume adjusted (to compensate for any differences in the loading and elution volume), where it can be loaded into the same or a different column. At each column step, the number of binding sites is adjusted to be equal to half of the total number of metal ions. Note that this decision is slightly different from the one we adopted in Medin *et al*.^8^ where the number of binding sites is adjusted so that half of all REE are biosorbed. Although these two options produce extremely similar results under the solution conditions and dissociation constants used in this article, they will diverge in when the REE concentration drops significantly below the dissociation constant. In an upcoming work we will explore different options for this decision. This process of load, bind, elute and re-load is repeated until the desired purity of M^1^ in the eluant is achieved (*f*_M1,b_ = 0.9, 0.95, or 0.99). While the effect of the selective biosorption on the purity of a target metal (say M_1_) is small in any individual step, the effect of successive enrichment is not. Numerical models of this process are implemented in the Python code ree-purification^41^.

The prospects of achieving high lanthanide purity by repeated biosorption and elution depend on the nature of the binding sites on the biosorption microbe’s surface. As of the time of writing, we can not uniquely specify all of the molecular details of lanthanide binding. As a result, we cannot perfectly predict the behavior of a repeated biosorption and elution process using the microbe. However, we can fit the preferences seen in our earlier work^8^ to two useful models (a single-site and a triple-site model), and then calculate the future behavior as the solution is enriched. For these examples, we have chosen Eu as the target metal for purification as wild-type *S. oneidensis* has a preference for it under low ionic strength, high lanthanide concentration^8^.

In the first model, the microbe uses a single type of binding site that is described by three dissociation constants (*e*.*g*., for Eu, Yb and La). A mathematical description of this model is included in **Note S2**. The preferences of the microbe for individual lanthanides are tuned by changing the relative values of the dissociation constants (**Figures 2A** and **2B**).

**Figure 2.**
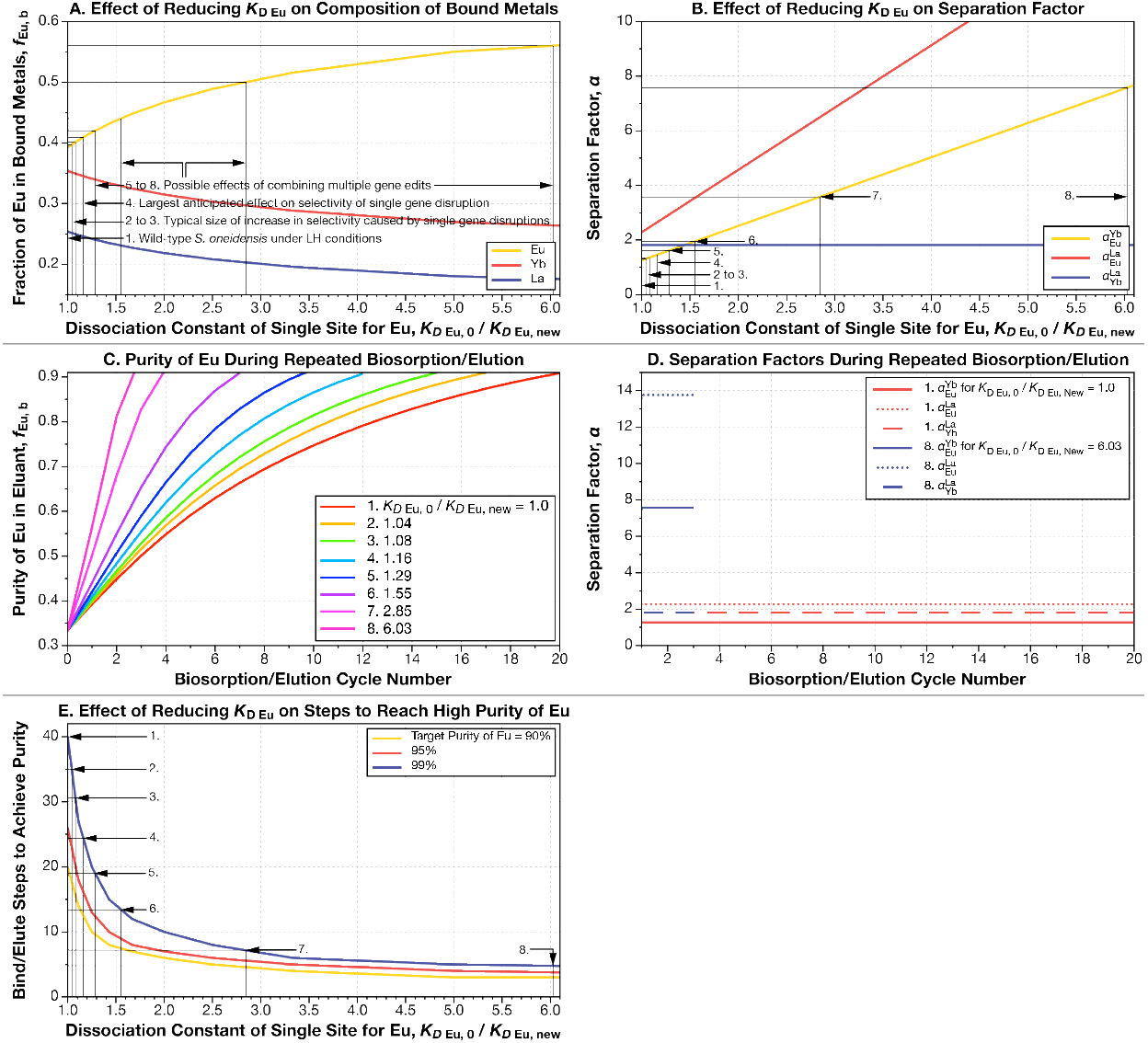
Lowering the dissociation constant of the single site, single microbe model for M_1_ (Eu) dramatically reduces the number of biosorption/elution cycles needed to achieve high purity. We fitted data on the fractions of each lanthanide (La, Eu, and Yb) bound by the wild-type *S. oneidensis* under low ionic strength, high REE-concentration conditions to a single-binding-site model with the Fit_KD.py program in the ree-purification repository^41^. We model the effect of reducing the dissociation constant of this single site for just Eu, while keeping the dissociation constants for Yb and La the same. See **Figure 1A, Note S1**, and **Table 1** for a full description of this model. (**A**) Reducing the Eu-dissociation constant of the REE-binding site raises the fraction of Eu initially bound to the surface of *S. oneidensis*. Note that the *x*-axis shows the inverse of dissociation constant for Eu. We have marked several notable binding-fractions on the plot (scenarios 1 to 8 in **Table 2**). Wild-type behavior (scenario 1) is at the far left of the plot. For example, an increase in the fraction of Eu bound to *S. oneidensis* by 1% from 39.3 to 39.7%, corresponds to a reduction in the dissociation constant by a factor 1.04 (scenario 2). An increase in *f*_Eu, b_ to 40.9% (a 4% increase over wild-type) corresponds to a reduction in *𝒦*_D Eu_ by a factor of 1.16 (scenario 4). (**B**) Reducing the dissociation constant for Eu raises the Eu-Yb and Eu-La separation factors, but does not affect the Yb-La separation factor. (**C**) Reducing the dissociation constant for Eu decreases the number of biosorption/elution cycles needed to reach high purity. Each curve corresponds to one of the example dissociation constants highlighted in panels **A** and **B** and **Table 2**. (**D**) The separation factors for Eu and Yb, Eu and La, and Yb and La all remain constant throughout the separation process. We have highlighted the highest and lowest example dissociation constants (scenarios 1 and 8) from panel **A**. Note that for clarity, the colors in panel **D** do not correspond to the colors in panel **C**. (**E**) Even small reductions in the dissociation constant of the binding site for Eu make large reductions in the number of biosorption/desorption cycles needed to achieve high purity. For example, if the dissociation constant is lowered by a factor of just 1.04 (scenario 2), the number of cycles needed to reach 99% purity drops from 40 cycles to 36. If the dissociation constant is lowed by a factor of 1.08 (scenario 3), then the number of cycles drops to 31. Panels in this figure can be reproduced with the programs Fig-2A-B.PY, Fig-2C-D.PY, and Fig-2E.py in the ree-purification repository^41^.

In the second model, the microbe uses three binding sites. The model contains a Eu-preferring site, a Yb-site, and a La-site. A mathematical description of this model is included in **Note S3**. In this model, we keep the dissociation constants of each site (one for each lanthanide) constant, and we tune the overall lanthanide preferences of the microbe by changing the population-fractions of the three sites (**Figures 3A** and **3B**).

**Figure 3.**
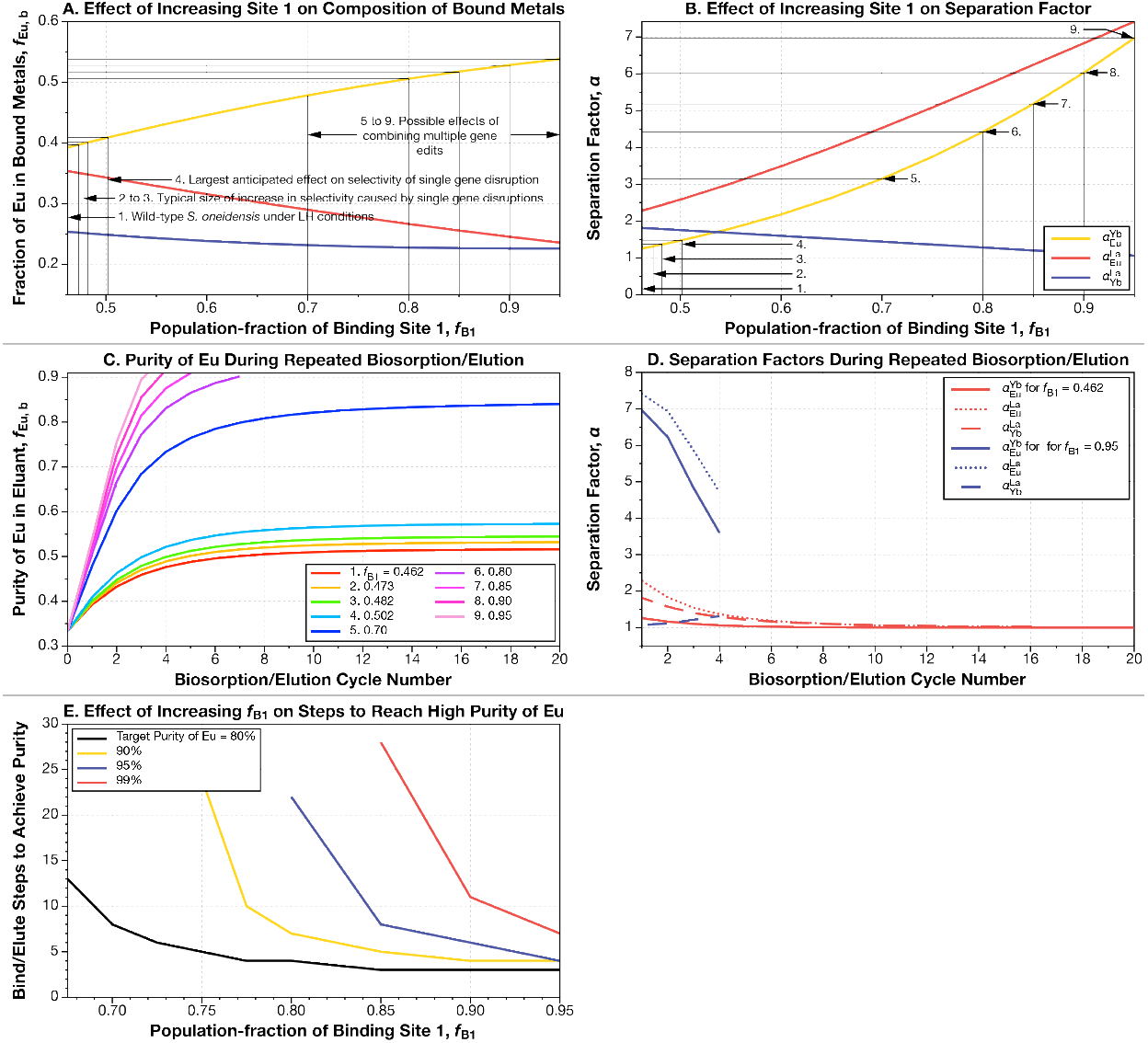
Raising the population-fraction of the Eu-preferring site (site 1) on the microbe’s surface increases the initial fraction of Eu bound, but does not allow high purity to be achieved until the population-fraction is raised above 90%. We fitted data on the fractions of each lanthanide (La, Eu, and Yb) bound by the wild-type *S. oneidensis* under low ionic strength, high REE-concentration conditions to a three-binding-site model with the Fit_fB_3S.py program. The model contains three binding sites, each with a fixed preference for one of the three REE in solution. The fit to the observed binding fractions of *S. oneidensis* is achieved by adjusting the population ratios of these sites. See **Figure 1A, Note S2**, and **Table 1** for full description of this model. (**A**) Raising the population of the Eu-preferring site (site 1) increases the initial fraction of Eu bound to the microbe’s surface. We have marked several notable binding-fractions on the plot (see **Table S5**). For example, an increase in the fraction of Eu bound to the microbe by 1% from 39.3 to 39.7% (scenario 2), corresponds to an increase in the population of the Eu-site from 46.2 to 47.3%. An increase in *f*_Eu, b_ to 40.9% (a 4% increase over wild-type; scenario 4) corresponds to an increase in the population of the Eu-site to 50.2%. (**B**) All separation factors change as the population-fractions of the three metal-binding sites change. (**C**) Increasing the population-fraction of the Eu-preferring site decreases the number of biosorption/ desorption cycles needed to increase Eu-purity However, unlike in the single-site model (**Figure 2**), most of the curves plateau well before reaching high purity. Each curve corresponds to one of the example dissociation constants highlighted in panels **A** and **B** and summarized in **Table 3**. (**D**) Unlike in the single-site model, the Eu-Yb, Eu-La, and Yb-La separation factors do not remain constant throughout the separation process. We have highlighted the highest and lowest example dissociation constants (*f*_b Eu, 39.3%_, and *f*_b Eu, 53%_) from panel **A**. (**E**) High purity of Eu is possible, but increasing the population fraction of the Eu-preferring site above 90% is necessary to do it. Panels in this figure can be reproduced with the programs Fig-3A-B.PY, Fig-3C-D.PY, and Fig-3E.py in the ree-purification repository^41^.

We can then ask what happens in a separation process if we change the parameters of the models to predict the effect of the small selectivity changes conferred by single gene modifications^8^. Finally, we can ask what happens if we further change the parameters of the models to predict the effects of more extensive genomic editing.

## Results and Discussion

While the effect of the selective biosorption on the purity of a target metal (*e*.*g*., Eu) is small in any individual step (**Figures 2A** and **3A**), the effect of successive enrichment is not. Furthermore, even small changes to binding selectivity make large changes to the length of separation process. The number of load-bind-elute cycles needed to achieve high purities are shown in **Figures 2C, 2E, 3C, 3E, 4A**, and **4B**.

**Figure 4.**
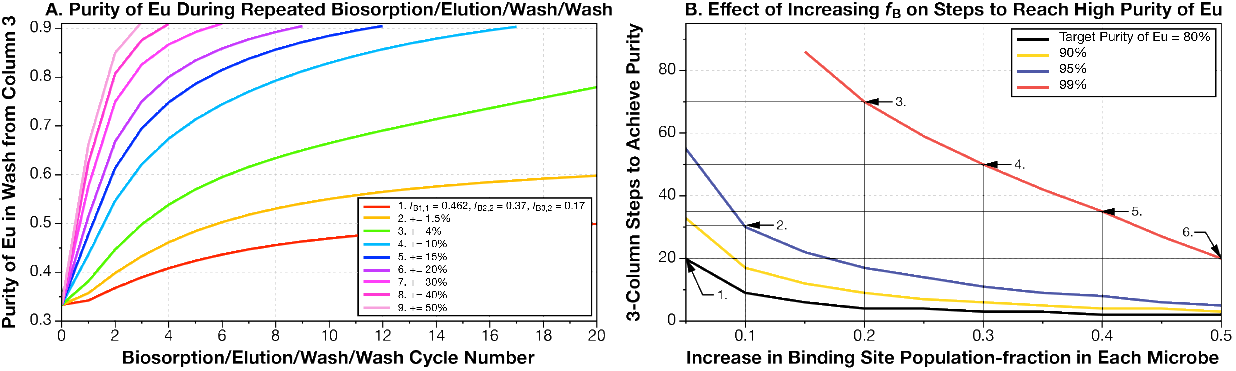
High purity of Eu can be achieved without large changes to the population-faction of the Eu-preferring site if a three-column, three-microbe system is used. As we noted in **Figures 3C** and **3E**, in contrast to using a microbe with a single type of binding site, repeated biosorption and desorption using a microbe with three binding sites does not always produce increasing purity of Eu, unless the population-fraction of the Eu-site is very high. To solve this problem, we turned to system where three columns each containing a different type of microbe is used in series (see **Figure 1B**). The first separates Eu, while the second 2 steps remove Yb and La respectively. Using this system with the wild-type microbes increases the purity of Eu slowly, but even modest changes to the population-fractions of one binding site in each of the microbes produces very notable reductions in steps to high purity. (**A**) The purity of Eu increases faster as we increase the population-fraction of the target site on each microbe. For example += 1.5% means that we increment the population of the Eu-site on the first microbe by 1.5% (*e*.*g*., from 0.462 to 0.477), increase the population of the Yb-site on the second microbe by 1.5% (*e*.*g*., from 0.370 to 0.385), and increase the population-fraction of the La-site on the third microbe by 1.5% (*e*.*g*., from 0.168 to 0.183). (**B**) Increasing the population-fraction of each of the three sites in each of the three microbes produces rapid improvements in the number of steps to high purity. Panels in this figure can be reproduced with the programs Fig-4A.py and Fig-4B.py in the ree-purification repository^41^.

### Fitting Single-site-type Binding Model to Experimental Data

We used a minimization algorithm included in the SCIPY package^42^ to fit two dissociation constants to the binding data for the wild-type *S. oneidensis*^8^ (note, we estimate that *𝒦*_D, Eu_ ≈ 10^-6^ M). This program can be found in Fit_KD.py script in the ree-purification repository^41^. We denote Eu, the metal with the highest concentration in the bound phase, as M_1_; Yb, the second highest concentration in bound phase, as M_2_; and La, the lowest concentration, as M_3_. We find, that relative to the dissociation constant for Eu (M_1_),

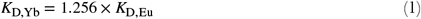

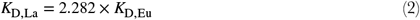

Thus, for a single-site system where the separation factors depend only upon the relative values of dissociation constants,

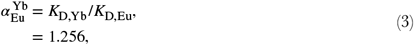

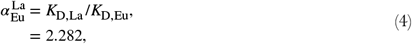

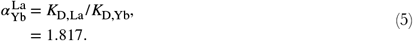

With estimates for the apparent relative dissociation constants of Eu, Yb and La, we can model the effects of improving the specificity of *S. oneidensis* for one of the metals. In **Figure 2A**, we reduce the dissociation constant of the binding site for Eu to 1/6^th^ of its original value, while holding its values constant for Yb and La. Reducing the dissociation constant of the site for Eu increases its binding fraction. In **Figure 2A**, we have marked several example binding fractions of Eu and their corresponding apparent dissociation constant for Eu (summarized in **Table 1**). For example, if the binding fraction of Eu rises from its original value 39.3% by 1% (the size of some of the smaller selectivity changes seen in Medin *et al*.^8^) to 39.7%, this corresponds to a decrease in the dissociation constant of the binding for Eu to 0.96 × its original value (or *𝒦*_D Eu, 0_ / *𝒦*_*D* Eu, New_ = 1.010). An increase in the binding fraction of Eu to 40.9% (a 4.1% increase, corresponding to one of the larger increases seen in our upcoming work by Medin *et al*.^8^), decreases the dissociation constant for Eu to 0.86 × its original value (or *𝒦*_D Eu, 0_ / *𝒦*_*D* Eu, New_ = 1.163). If we could increase the binding fraction of Eu to 56%, this would correspond to a decrease in the dissociation constant for Eu to 0.17 × its original value (or *𝒦*_D Eu, 0_ / *𝒦*_*D* Eu, New_ = 6.027). The corresponding separation factors for Eu and Yb; Eu and La; and Yb and La are shown in **Figure 2B**. As should be expected, the separation factors for Eu and Yb, and Eu and La increase as the dissociation constant for Eu is reduced, but the separation factor for Yb and La remains constant.

### Effect of Improvement of Single-site Binding Specificity on REE-separations

We used the dissociation constants for Eu, Yb, and La (**Equations 1** and **2**) to calculate the separation behavior of the single-site model. This model is always able to achieve arbitrarily high purity (**Figures 2C** to **2E**). If the single-site model representing wild-type *S. oneidensis* repeatedly binds and elutes an initially equimolar solution of Eu, Yb, and La, it would take 20 cycles to reach 90% purity of Eu; 26 cycles for 95% purity; and 40 cycles for 99% purity (**Figure 2E** and **Table 2**).

In the single-site model, even mutants that make small changes to the initial binding fractions of metals make big changes to separation process length (**Figures 2C** to **2E, Table 2**). If we increase the initial binding fraction of Eu by 1% (from 39.3% to 39.7%; a fairly typical increase produced for example by disruption of the *wzz* gene^8^) this corresponds to a 1.04-fold reduction in the Eu-dissociation constant (**Scenario 2** in **Figure 2A**). If this mutant were used in repeated enrichment, then only 36 biosorb/elute cycles are needed for 99% purity (**Figures 2C** and **2E, Table 1**). A 2% increase in initial binding fraction (*e*.*g*, comparable to disruption of the *wbpA* gene^8^) drops the steps to 99% Eu-purity to 31. If we could achieve a 4% increase in initial preference for Eu (the largest size of change we see due to single-gene disruptions) then we could reduce the process length to 25 steps. Increasing the initial binding preference to 50% Eu (*e*.*g*., by combining mutations to multiple genomic loci in a single strain), then we could reduce the steps to 99% purity to only 8 (**Figures 2C** and **2E, Table 1**)

### Fitting Three-site-type Binding Model to Experimental Data

We modeled the selectivity of *S. oneidensis* by changing the fraction of the total binding sites made up by each of these three binding sites and by changing the dissociation constant of each site for its target metal. To perform this fit, we used a minimization algorithm using the SCIPY package to fit two dissociation constants to the binding data for the wild-type *S. oneidensis* from our recent work^8^. This algorithm can be found in the Fit_fB_3S.py in the ree-purification repository^41^.

The first site has a preference for Eu (note that the choice of 8 was somewhat arbitrary, but it does allow us to have similarly sized population-fractions for sites 1, 2, and 3) and is consistent with our single-site model,

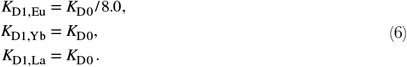

The number of site 1 molecules, as a fraction of the total,

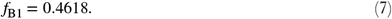

The second site has a preference for Yb,

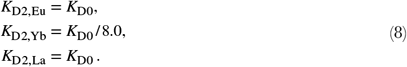

The number of site 2 molecules, as a fraction of the total,

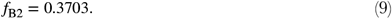

Finally, the third site has a preference for La,

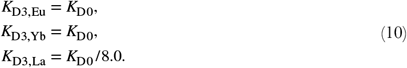

The number of site 3 molecules, as a fraction of the total,

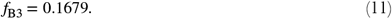

Although the separation factors in the three-site model do not result from simple ratios of dissociation constants (**Note S3**), the overall numerical results is the same as in **Equations 3** to **5**.

Intuitively, we would expect that as we increase the fraction of binding site 1 (the site with a preference for Eu), the fraction of Eu bound to *S. oneidensis* should increase. In **Figure 3A**, we increase the fraction of site 1 from its original value (46.3%) to 95%, while keeping all dissociation constants the same. As the fraction of site 1 increases, the fractions of sites 2 and 3 are reduced, but maintain their original size ratio. For example, in the original scenario that mimics the wild-type behavior, sites 2 and 3 occupy 53.7% of the total sites (100 – 46.3%). Thus, site 2 occupies 69% of the remaining sites, and site 3 occupies 31% of the remainder. As the fraction of 1 increases, the fraction of sites 2 and 3 in the remainder remains constant. So, when *f*_B1_ = 95%, *f*_B2_ = 3.4% (69% of 5%), while *f*_B3_ = 1.6% (31% of 5%).

In **Figure 3A**, we have marked several example binding fractions of Eu (see **Table 3** for a summary) and the corresponding calculated population-fraction of site 1 (the site with a preference for Eu). For example, if the binding fraction of Eu (*f*_Eu, b_) rises from its original value 39.3% by 1% (the size of some of the smaller selectivity changes seen in Medin *et al*.^8^) to 39.7%, this corresponds to an increase in the population of binding site 1 to 47.3%.

An increase in the binding fraction of Eu to 40.9% (a 4% increase, corresponding to one of the larger increases in Medin *et al*.^8^), corresponds to an increase in the population of binding site 1 to 48.2%. The corresponding separation factors for Eu and Yb; Eu and La; and Yb and La are shown in **Figure 3B**. Unlike in **Figure 3B**, the separation factor for Yb and La does not remain constant.

### Effect of Improvement in Three-site Model on REE-separations

What does the increase in the fraction of binding site 1 (the site with a preference for Eu) mean for the purification of Eu? If we repeatedly pass the eluant of the biosorption column through it, we should expect to enrich the solution for Eu (as shown in **Figure 1A**). A simulation of this process is shown in **Figure 3C**. As the eluant solution is passed through the column, the purity of Eu does increase. However, unlike in **Figure 2C** (where the purity of Eu does always eventually reach the target of 90%), for lower fractions of site 1, the purity of Eu almost plateaus. This is because in the three-site model, unless you can completely eliminate them, there are always places on the surface of the microbe for Yb and La to strongly bind. Unlike in the case where we have a single binding site type, where any change to the microbe is useful and reduces the steps to get to 99% purity, when there are multiple sites that are strongly selective for a single lanthanide, you need to eliminate a very high fraction of them to make a useful microbe. For example, even if the fraction of site 1 is raised from 46.3 to 80%, we will require 22 bind/ elute cycles to reach 95% purity (**Figure 3E, Table 3**).

In the three-site model, extensive genetic editing will be needed to achieve high purity of Eu. Under the single-site model, it takes the wild-type 40 bind/elute cycles to reach 99% Eu-purity (**Figure S9D** and **Table S4**).

### Three Microbe Separation System

The key result of our upcoming work is that there are few if any, dominant players in REE-biosorption in *S. oneidensis*. Instead of one dominant type of binding site like teichoic acids in *Bacillus subtilis*, many different types of sites, encoding structural features ranging from pilus to lipopolysaccharides to outer membrane proteins, contribute to the adsorption of REE to the *S. oneidensis* membrane. This result suggests that in a successive enrichment process *S. oneidensis* has the potential to behave like the three-site model (**Note S3, Figure 3**), and might produce plateauing purity before an industrially-acceptable level is reached.

However, even if *S. oneidensis* does produce plateauing purity in a repeated enrichment process, our model of REE-separation indicates that this problem is not insurmountable. The model suggests that a process that uses multiple microbes in series (**Figures 1B** and **4**; one to enrich for the target lanthanide, and others to remove the competing lanthanides) could achieve industrially-acceptable purity with only small genetic edits.

When used in three-microbe scheme, the three-site model representing wild-type *S. oneidensis* is only able to achieve ≈ 50% Eu-purity (**Figure 4A**). However, if the population-fractions of the target binding sites on each microbe are incremented by just 1.5% then performance improves dramatically (*i*.*e*., the population of the Eu-preferring site on microbe 1 is increased from 46.2 to 47.7%; the fraction of the Yb-site on microbe 2 from 37.0 to 38.5%; and the La-site in microbe 3 from 16.8 to 18.3%). If we increment the population fraction of the target sites by 4% in each microbe (scenario 3 in **Figure 4A**), the purity of Eu starts to take-off. Notably, these increases in population-fraction are typical of single genomic loci mutations (see **Table 2** for connection).

The three-microbe system allows us to reach high purity of Eu by repeated biosorption and elution without heroic genetic engineering efforts. In order to reach 99% purity, we estimate that population-fraction of the important site in each microbe must be incremented by ≈ 15%. The number of steps (in this case passes through the 3 microbe system) to reach 90, 95, and 99% purity are shown in **Figure 4** and **Table 4**. This suggests that this three microbe column system could be made to operate by combining together microbes with edits to only one or two genetic loci.

**Table 4.** Summary changes to REE-biosorption selectivity on REE-separation with a three-microbe, three-site-type binding model. This table summarizes the results of **Figure 4**. (**1**) Entries correspond to marked points on **Figure S9**. (**2**) Increase in the population-fraction of the targeted metal-preferring site on each microbe. For example += 5% means that we increment the population of the Eu-site on the first microbe by 5% (*e*.*g*., from 0.462 to 0.512), increase the population of the Yb-site on the second microbe by 5% (*e*.*g*., from 0.370 to 0.420), and increase the population-fraction of the La-site on the third microbe by 5% (*e*.*g*., from 0.168 to 0.218). (**3** to **6**). Reduction in biosorption/desorption cycles needed to achieve a Eu purity of 80, 90, 95, and 99%. This table can be reproduced with the programs Fig-4A.PY, and Fig-4B.py in the ree-purification repository^41^.

| 1. Scenario | 2. Increase in Population of Primary Binding Site | 3. 80% Purity Eu | 4. 90% Purity Eu | 5. 95% Purity Eu | 6. 99% Purity Eu |
| --- | --- | --- | --- | --- | --- |
| 1 | 0.05 | 20 | 33 | 55 |  |
| 2 | 0.10 | 9 | 17 | 30 |  |
| 3 | 0.20 | 4 | 9 | 17 | 70 |
| 4 | 0.30 | 3 | 6 | 11 | 50 |
| 5 | 0.40 | 2 | 4 | 8 | 35 |
| 6 | 0.50 | 2 | 4 | 6 | 21 |

## Conclusions

In this article, we present three models of lanthanide separation by repeated biosorption and desorption. The first model describes a microbe with a single type of binding site (most applicable to immobilized chelators), while the second describes a microbe with three slightly more selective binding sites. The final model uses three microbes in series, the first to bind the target metal, and the second and third to remove contaminants.

Despite the small size of changes to biosorption preference (≈ 1 to 4%) caused by disruptions to single genomic loci observed in our recent work^8^, these changes might produce large reductions in the length of a repeated enrichment process for individual lanthanides (**Figure 2** and **Table 2**). If *S. oneidensis* behaves like it has a single type of binding site (this does not mean that *S. oneidensis* only has one physical species of binding site, only that its separation factors for multiple lanthanides remains constant as the external concentrations change (**Note S2** and **Figure 2B**), then even the mutants that we have right now^8^ could be used to reduce the length of the separation process for Eu by up to ≈ 25% (**Note S2, Figure 2E**, and **Table 2**). If edits to multiple genomic loci could be combined in a single strain, then it is possible that the length of the purification process could be reduced even more.

On the other hand, if a microbe has multiple binding sites (each with a relatively strong preference for individual lanthanides, *e*.*g*., a separation factor of ≈ 8), then purification by repeated biosorption and elution becomes difficult without large genetic engineering efforts (**Figures 3C** and **3D**).

Even though small edits to a single microbe with multiple strongly selective sites will not be useful in purifying an individual lanthanide, we hypothesized that by combining multiple microbes together in a purification process we could achieve high purity (**Figure 4**). This system enriches for M_1_ in the first column (*e*.*g*., Eu); and then removes M_2_ and M_3_ in the second and third columns respectively. This system allows us to reach high purity of Eu by repeated biosorption and elution without heroic genetic engineering efforts. In order to reach 99% purity, we estimate that population-fraction of the important site in each microbe must be incremented by ≈ 15% (site 1 in microbe 1 from 0.462 to 0.612; site 2 in microbe 2 to 0.52; and site 3 in microbe 3 to 0.318). Taken together, these results give a roadmap for building, operating, and troubleshooting biosorption-based rare earth separation systems.

## End Notes

### Code Availability

All code used in calculations in this article is available at https://github.com/barstowlab/ree-purification and is archived on Zenodo^41^.

### Materials & Correspondence

Correspondence and material requests should be addressed to B.B.

### Author Contributions

Conceptualization, C.A., S.M. and B.B.; Methodology, C.A., S.M. and B.B.; Investigation, C.A., S.M., J.A., B.D., E.E., C.K., J.L., T.J.S., D.S., Z.T. M.X., K.Z., B.B; Writing - Original Draft, J.A., B.D., E.E., C.K., J.L., T.J.S., D.S., Z.T. M.X., K.Z., B.B; Writing - Review & Editing, B.B.; Funding Acquisition, B.B.; Resources, M.R., E.G., M.W., and B.B.; Supervision, B.B.; Data Curation, B.B.; Visualization, B.B..

## Acknowledgements

S.M. was supported by a Cornell Presidential Life Sciences Graduate Fellowship. This work was supported by Cornell University startup funds, an Academic Venture Fund award from the Atkinson Center for Sustainability at Cornell University, a Career Award at the Scientific Interface from the Burroughs Welcome Fund to B.B., ARPA-E award DE-AR0001341 to B.B.., Air Force Research Laboratory award FA8750-22-2-0500 to B.B., and a gift from Mary Fernando Conrad and Tony Conrad to B.B..

