## Supplementary Notes and Figures for "Constraints on Lanthanide Separation by Selective Biosorption"

#### Supplementary Information Tables

**Table S1.** Symbols used in this article.

**Table S2.** Representative values of metal biosorption per unit mass of dry biomass.

#### Supplementary Information Notes

**Note S1.** Model for separation system using a single type of binding site.

**Note S2.** Model for separation system using three types of binding site.

**Note S3.** Comparison of experimental data with literature observations of biosorption

| Symbol | Unit | Description |
|---|---|---|
| $a_x^y$ | # | Separation factor for metals $x$ and $y$. $a_x^y = D_y / D_x$. |
| $c_{Bx,f}$ | Mol $L^{-1}$ | Concentration of binding site x that is unoccupied. |
| $c_{Mx,b}$ | Mol $L^{-1}$ | Concentration of metal $x$ in the bound phase (used in single-site model). |
| $c_{Mx,by}$ | Mol $L^{-1}$ | Concentration of metal $x$ bound to site $y$. |
| $c_{Mx,f}$ | Mol $L^{-1}$ | Concentration of metal $x$ in the solution phase. |
| $D_x$ | # | Distribution coefficient for metal $x$. The ratio of Mx in the free and bound phases. $D_x = c_{Mx,f} / c_{Mx,b}$. |
| $f_{Bx}$ | # | Fraction of total binding sites in immobilized biomass made up of site $x$. |
| $f_{Mx,b}$ | # | Fraction of metal $x$ (*e.g*, $M_1$ or Eu) bound to immobilized biomass. |
| $f_{Mx,f}$ | # | Fraction of metal $x$ in the free (solution) phase. |
| $K_{D,x}$ | Mol $L^{-1}$ | Dissociation constant of single binding site for $M_x$ (*e.g*, $K_{D,Eu}$). |
| $K_{Dx,y}$ | Mol $L^{-1}$ | Dissociation constant of binding site $x$ for $M_y$ (*e.g*, $K_{D1,Eu}$). |
| $N_A$ | molecule $mol^{-1}$ | Avogadro constant. |
| $n_{B,T}$ | Mol | Number of binding sites on biosorption column. |
| $n_{MT}$ | Mol | Total number of metal ions. |
| $n_{Mx,T}$ | Mol | Number of metal $x$ (*e.g*, Eu) ions loaded into separation column. |
| $V_{load}$ | L | Volume of solution loaded into biosorption column. |

**Table S1.** Symbols list.

| **Organism** | **Loading Capacity (mg per g dry weight)** | **Metal** | **Reference** |
|---|---|---|---|
| *Bacillus circulans* | 5.8 to 26.5 | Cd | [Yilmaz2005a] |
| *Saccharomyces cerevisiae* | 42 to 60 | $Ag^{+}$ | [Simmons1996a] |
| *Spirogyra species* | 133.3 | $Cu^{2+}$ | [Gupta2006a] |
| *Talaromyces emersonii* | 280 | $U^{6+}$ | [Bengtsson1995a] |
| *Aspergillus niger* | 31.9 to 97.6 | $Zn^{2+}$ | [Luef1990a] |
| *Penicillium chrysogenum* | 19.9 to 85.5 | $Zn^{2+}$ | [Luef1990a] |
| *Claviceps paspali* | 31.9 to 97.6 | $Zn^{2+}$ | [Luef1990a] |
| *Saccharomyces cerevisiae* | 59 | Ag | [Bustard1998a] |
| *Saccharomyces cerevisiae* | 189 | Pb | [Bustard1998a] |
| *Citrobacter freundii* | 7.2 to 35.8 | Pb | [AlGarni2005a] |
| *Klebsiella pneumoniae* | 6.2 to 31.9 | Pb | [AlGarni2005a] |

**Table S2.** Representative values of metal biosorption per unit mass of dry biomass.

### Supplementary Information Notes

#### Note S1. Comparison of Experimental Data with Literature Observations of Biosorption

We fit data shown in Medin *et al.* [Medin2023a] on biosorption by wild-type *S. oneidensis* to the single- and triple-site models of binding (**Notes S2** and **S3**) to establish baseline parameters for our predictions of separation behavior by repeated binding and de-binding.

To sanity check the connection between experimental data and our models, we first calculate the lanthanide binding per gram of dry weight of *S. oneidensis* and compare with literature values. For the purposes of this analysis, we will restrict discussion to the low ionic strength, high REE conditions (LH) because these have the highest binding of REE, so we have to worry least about interference from NaCl.

Under the low ionic strength, high REE conditions (LH), the true wild-type *S. oneidensis* binds a total of 35.7 nanomoles (nmol) of lanthanides, from a total of 72 nmol in solution. The volume in these experiments is 400 μL, with an optical density of 0.85. Assuming that there are approximately $10^9$ cells per OD per mL of culture, then in each REE-binding experiment there are a total of $3.4 \times 10^8$ cells. This corresponds to $6.3 \times 10^7$ atoms of lanthanide per cell.

How does the mass of lanthanides bound per unit of dry weight compare with literature values? Under LH conditions *S. oneidensis* binds 8.9 nmol of La (25.4% of the total); 12.4 nmol Yb (35.3%); and 13.8 nmol of Eu (39.3%). These correspond to 3.6 fg La per cell; 6.3 fg Yb per cell; and 6.1 fg Eu per cell. This corresponds to a total of 16.1 fg total lanthanides per cell.

What is the dry weight per cell of *S. oneidensis*? We assume that *S. oneidensis* has a similar dry weight to *E. coli* which ranges 280 fg under 40 minute doubling conditions (BioNumbers ID (BNID) 103904); 480 fg (with a range of 358 to 622; BNID 102230); to 640 fg (BNID 100009). The average of these values is 470 fg. Thus the approximate loading capacity of *S. oneidensis* is s 16.1 fg $cell^{-1}$ / 470 fg dry weight $cell^{-1} \approx 30$ mg $g^{-1}$ of dry weight. This number is compares very well to literature values of metal binding per unit mass of dry biomass assembled in **Table S2**.

#### Note S2. Model for Separation System Using a Single Type of Binding Site

In the simplest model of the lanthanide separation scheme, we consider biomass that contains a single type of binding site (results shown in **Figure 2** and **Table 2** in the main text). The binding site has dissociation constants $K_{D,1}$, $K_{D,2}$, and $K_{D,3}$ for $M_1$, $M_2$, and $M_3$ respectively. After equilibration, the bound and free concentrations of $M_1$ ($c_{M1,f}$, $c_{M1,b}$) and the concentration of free binding sites ($c_{B,f}$) are related by dissociation constant $K_{D,1}$,

$$K_{D,1} = \left(c_{M1,f}\, c_{B,f}\right)/c_{M1,b}\,. \tag{S1}$$

Likewise, for metals 2 and 3,

$$K_{D,2} = \left(c_{M2,f}\, c_{B,f}\right)/c_{M2,b}, \tag{S2}$$

$$K_{D,3} = \left(c_{M3,f}\, c_{B,f}\right)/c_{M3,b}. \tag{S3}$$

The number of moles of each metal atom in the bound and free phases is just the concentration multiplied by the system volume. For example,

$$n_{M1,f} = c_{M1,f}\, V_{load}, \tag{S4}$$

$$n_{M1,b} = c_{M1,b}\, V_{load}, \tag{S5}$$

The total number of binding sites is equal to the number of free and bound sites,

$$n_{B,T} = V_{load}\left(c_{B1,f} + c_{B1,b}\right). \tag{S6}$$

The total number of each metal is just the sum of bound and free ions,

$$n_{M1,T} = V_{load}\left(c_{M1,f} + c_{M1,b}\right), \tag{S7}$$

$$n_{M2,T} = V_{load}\left(c_{M2,f} + c_{M2,b}\right), \tag{S8}$$

$$n_{M3,T} = V_{load}\left(c_{M3,f} + c_{M3,b}\right)\,. \tag{S9}$$

Parameters for the solution of this system of 8 equations were managed with a custom code (the function BINDSTATE_3METALS_1SITE in CONCENTRATIONSOLVERUTILS8 [Barstow2023a]) written in PYTHON, and the equations were solved numerically using SYMPY [Meurer2017a]. After the solution to the equations are found, the system is re-started, using just the bound metal ions as the new total metal ions for the next round ($n_{M1,b}$, $n_{M2,b}$, and $n_{M3,b}$).

The ratio of concentrations of each metal in the free and bound phases are called distribution coefficients,

$$D_1 = c_{M1,f}/c_{M1,b}, \tag{S10}$$

$$D_2 = c_{M2,f}/c_{M2,b}, \tag{S11}$$

$$D_3 = c_{M3,f}/c_{M3,b}\,. \tag{S12}$$

Furthermore, the ratio of distribution coefficients between metals are called separation factors,

$$\alpha_1^2 = D_2/D_1, \tag{S13}$$

$$\alpha_1^3 = D_3/D_1, \tag{S14}$$

$$\alpha_2^3 = D_3 / D_2 \,. \tag{S15}$$

For a system containing only a single type of binding site (or at least binding sites that can all be described by a single set of dissociation constants) the separation factors can be expressed simply. For example,

$$\begin{aligned} \alpha_1^2 &= D_2 / D_1, \\ &= \left(c_{\mathrm{M2,f}} / c_{\mathrm{M2,b}}\right) / \left(c_{\mathrm{M1,f}} / c_{\mathrm{M1,b}}\right), \\ &= \left(K_{\mathrm{D,2}} / c_{\mathrm{B,f}}\right) / \left(K_{\mathrm{D,1}} / c_{\mathrm{B,f}}\right) \\ &= K_{\mathrm{D,2}} / K_{\mathrm{D,1}} \,. \end{aligned} \tag{S16}$$

Likewise, when there is only one type of binding site,

$$\alpha_1^3 = K_{\mathrm{D,3}} / K_{\mathrm{D,1}}, \tag{S17}$$

$$\alpha_3^2 = K_{\mathrm{D,2}} / K_{\mathrm{D,3}} \,. \tag{S18}$$

As a result of this, the separation behavior of the system depends on the ratio of dissociation constants rather than their absolute magnitude. Furthermore, as the separation factor depends upon intrinsic molecular properties, it remains constant as the concentrations of metals and binding sites change (see **Figure 2D** for an example of this, and **Figure 3D** for an example of where it does not).

As the separation process proceeds, the number of binding sites in the column is adjusted to be equal to half the total number of ions in the load solution (this could be accomplished by sliding the biosorption material out of the loaded solution). Thus,

$$n_{\mathrm{B,T}} = 0.5 \times n_{\mathrm{MT}} \tag{S19}$$

Finally, to compare with experiment, we calculate the free and bound fractions of each metal in each phase. The proportion of each metal in the free (liquid) fraction,

$$f_{\mathrm{M1,f}} = n_{\mathrm{M1,f}} / \left(n_{\mathrm{M1,f}} + n_{\mathrm{M2,f}} + n_{\mathrm{M3,f}}\right), \tag{S40}$$

$$f_{\mathrm{M2,f}} = n_{\mathrm{M2,f}} / \left(n_{\mathrm{M1,f}} + n_{\mathrm{M2,f}} + n_{\mathrm{M3,f}}\right), \tag{S41}$$

$$f_{\mathrm{M3,f}} = n_{\mathrm{M3,f}} / \left(n_{\mathrm{M1,f}} + n_{\mathrm{M2,f}} + n_{\mathrm{M3,f}}\right) . \tag{S42}$$

and in the bound fractions,

$$f_{\mathrm{M1,b}} = n_{\mathrm{M1,b}} / \left(n_{\mathrm{M1,b}} + n_{\mathrm{M2,b}} + n_{\mathrm{M3,b}}\right), \tag{S43}$$

$$f_{\mathrm{M2,b}} = n_{\mathrm{M2,b}} / \left(n_{\mathrm{M1,b}} + n_{\mathrm{M2,b}} + n_{\mathrm{M3,b}}\right), \tag{S44}$$

$$f_{\mathrm{M3,b}} = n_{\mathrm{M3,b}} / \left(n_{\mathrm{M1,b}} + n_{\mathrm{M2,b}} + n_{\mathrm{M3,b}}\right) . \tag{S45}$$

We fit data on the fractions of Eu [Medin2023a] ($f_{\mathrm{M1,b}}$ or $f_{\mathrm{Eu,\,b}}$); Yb ($f_{\mathrm{M2,b}}$ or $f_{\mathrm{Yb,\,b}}$); and La ($f_{\mathrm{M3,b}}$ or $f_{\mathrm{La,\,b}}$) bound to wild-type *S. oneidensis* to initialize our model.

#### Note S3. Model for Separation System Using Three Types of Binding Site

A more complex model of lanthanide separation considers multiple types of binding sites. Here we consider a system that contains three types of binding site, each capable of binding all of the three metals in the system (results shown in **Figure 3** and **Table 3** in the main text). This system is described by 18 simultaneous equations (in contrast to the 8 equations needed to describe a system with one binding site).

The dissociation constant of site 1 for $M_1$,

$$K_{D1,1} = (c_{M1,f}\, c_{B1,f})/c_{M1,b1}\,. \tag{S20}$$

where $c_{B1,f}$ is the concentration of free type 1 sites, and $c_{M1,b1}$ is the concentration of site 1 molecules bound to $M_1$. Likewise, the dissociation constant of site 1 for $M_2$,

$$K_{D1,2} = (c_{M2,f}\, c_{B1,f})/c_{M2,b1}\,. \tag{S21}$$

where $c_{M2,b1}$ is the concentration of site 1 molecules bound to $M_2$. Likewise,

$$K_{D1,3} = (c_{M3,f}\, c_{B1,f})/c_{M3,b1}\,. \tag{S22}$$

where $c_{M3,b1}$ is the concentration of site 1 molecules bound to $M_3$. Furthermore, for sites 2 and 3,

$$K_{D2,1} = (c_{M1,f}\, c_{B2,f})/c_{M1,b2}, \tag{S23}$$

$$K_{D2,2} = (c_{M2,f}\, c_{B2,f})/c_{M2,b2}, \tag{S24}$$

$$K_{D2,3} = (c_{M3,f}\, c_{B2,f})/c_{M3,b2}, \tag{S25}$$

$$K_{D3,1} = (c_{M1,f}\, c_{B3,f})/c_{M1,b3}, \tag{S26}$$

$$K_{D3,2} = (c_{M2,f}\, c_{B3,f})/c_{M2,b3}, \tag{S27}$$

$$K_{D3,3} = (c_{M3,f}\, c_{B3,f})/c_{M3,b3}\,. \tag{S28}$$

The total number of metal 1, 2, and 3 ions,

$$n_{M1,T} = V_{load}\,(c_{M1,f} + c_{M1,b1} + c_{M1,b2} + c_{M1,b3}), \tag{S29}$$

$$n_{M2,T} = V_{load}\,(c_{M2,f} + c_{M2,b1} + c_{M2,b2} + c_{M2,b3}), \tag{S30}$$

$$n_{M3,T} = V_{load}\,(c_{M3,f} + c_{M3,b1} + c_{M3,b2} + c_{M3,b3})\,. \tag{S31}$$

The total number of binding sites of type 1, 2 and 3,

$$n_{B1,T} = V_{load}\,(c_{B1,f} + c_{M1,b1} + c_{M2,b1} + c_{M3,b1}), \tag{S32}$$

$$n_{B2,T} = V_{load}\,(c_{B2,f} + c_{M1,b2} + c_{M2,b2} + c_{M3,b2}), \tag{S33}$$

$$n_{B3,T} = V_{load}\,(c_{B3,f} + c_{M1,b3} + c_{M2,b3} + c_{M3,b3})\,. \tag{S34}$$

Finally, the concentration of bound sites of type 1, 2, and 3,

$$c_{B1,b} = c_{M1,b1} + c_{M2,b1} + c_{M3,b1}, \tag{S35}$$

$$c_{B2,b} = c_{M1,b2} + c_{M2,b2} + c_{M3,b2}, \tag{S36}$$

$$c_{\mathrm{B3,b}} = c_{\mathrm{M1,b3}} + c_{\mathrm{M2,b3}} + c_{\mathrm{M3,b3}}\,. \tag{S37}$$

The last three equations (**S35** to **S37**) are not strictly necessary but do help to stabilize the numerical solution.

Parameters for the solution of this system of 18 equations were managed with a custom code (the function BINDSTATE_3METALS_3SITES in CONCENTRATIONSOLVERUTILS8 [Barstow2023a]) written in PYTHON, and the equations were solved numerically using SYMPY [Meurer2017a].

The distribution coefficients can be calculated similarly to **Equations S10** to **S12**, but are complicated by the fact that each metal binds to more than one site. For example,

$$D_1 = \frac{c_{\mathrm{M1,f}}}{c_{\mathrm{M1,b1}} + c_{\mathrm{M1,b2}} + c_{\mathrm{M1,b3}}}\,. \tag{S38}$$

Substituting in **Equations S20** to **S28**,

$$\begin{aligned} D_1 &= \frac{c_{\mathrm{M1,f}}}{c_{\mathrm{M1,f}}/\left(c_{\mathrm{B1,f}}\,K_{\mathrm{D1,1}}\right) + c_{\mathrm{M1,f}}/\left(c_{\mathrm{B2,f}}\,K_{\mathrm{D2,1}}\right) + c_{\mathrm{M1,f}}/\left(c_{\mathrm{B3,f}}\,K_{\mathrm{D3,1}}\right)}\,, \\ &= \frac{1}{1/\left(c_{\mathrm{B1,f}}\,K_{\mathrm{D1,1}}\right) + 1/\left(c_{\mathrm{B2,f}}\,K_{\mathrm{D2,1}}\right) + 1/\left(c_{\mathrm{B3,f}}\,K_{\mathrm{D3,1}}\right)}\,. \end{aligned} \tag{S39}$$

As in **Note S1**, we compare our results to experiment by comparing the calculated free and bound fractions of each metal in each phase.
